# CellGO: A novel deep learning-based framework and webserver for cell type-specific gene function interpretation

**DOI:** 10.1101/2023.08.02.551654

**Authors:** Peilong Li, Junfeng Wei, Ying Zhu

**Affiliations:** State Key Laboratory of Medical Neurobiology, MOE Frontiers Center for Brain Science, Institutes of Brain Science and Department of Neurosurgery, Huashan Hospital, Fudan University, Shanghai 200032, China

**Author notes:** Correspondence to: Ying Zhu. These authors contributed equally to this work. Peilong Li, Junfeng Wei.

## Abstract

Interpreting the function of genes and gene sets identified from omics experiments remains a challenge, as current pathway analysis tools often fail to account for complex interactions across genes and pathways under specific tissues and cell types. We introduce CellGO, a tool for cell type-specific gene functional analysis. CellGO employs a deep learning model to simulate signaling propagation within a cell, enabling the development of a heuristic pathway activity measuring system to identify cell type-specific active pathways given a single gene or a gene set. It is featured with additional functions to uncover pathway communities and the most active genes within pathways to facilitate mechanistic interpretation. This study demonstrated that CellGO can effectively capture cell type-specific pathways even when working with mixed cell-type markers. CellGO’s performance was benchmarked using gene knockout datasets, and its implementation effectively infers the cell type-specific pathogenesis of risk genes associated with neurodevelopmental and neurodegenerative disorders, suggesting its potential in understanding complex polygenic diseases. CellGO is accessible through a python package and a four-mode web interface for interactive usage with pretrained models on 71 single-cell datasets from human and mouse fetal and postnatal brains.

## 1 Introduction

The prevalence of high-throughput technologies has greatly accelerated the discovery of genes with specific functions and their associations with diseases (1-5). For instance, a genome-wide association study (GWAS) can reveal hundreds of thousands of disease risk genes (6-13), and a single RNA-seq experiment may detect from one to thousands of differentially expressed genes (14-20). However, the challenge remains in mechanistic interpretation of these discoveries.

The common strategy to gain mechanistic insights from genes is pathway analysis (21,22), and varieties of tools have been developed and widely used. In spite of various forms, the underlying concept of these tools is to find a statistic to measure the pathway ‘activities’ given a set of genes. The pathway ‘activity’ can be assessed simply by quantifying the number of genes shared between the gene set for testing and the pathway. Subsequently, the pathways deemed ‘active’ are chosen if the proportion of overlapped genes in the tested gene set is significantly greater than that found in the background (23). This approach is called over-representation analysis (ORA) and is adopted by many popular tools (24-29). However, ORA relies on the assumption that all genes make independent and equal contributions to pathways, which may not always hold true in biological systems. To overcome this limitation, functional class scoring methods (FCS) have been proposed (30,31). FCS computes pathway activities by aggregating gene-level scores (32,33), such as expression fold changes and *P*-values, which reflect the contributions of genes to each pathway. However, neither ORA nor FCS takes into account the interrelationship between genes, which can also influence a gene’s contribution to the pathway activity. Pathway topology-based methods (34-38) (PTM) have been developed to address this limitation by integrating the interconnections between genes within pathways, potentially improving performance over previous approaches. Nevertheless, none of the three widely-used categories of methods considers crosstalk between pathways.

Moreover, the biological context in which pathways are active (e.g., specific tissues or cell types) can provide critical information for mechanistic explanations (22). Unfortunately, current methods often lack this consideration. Although a recent study proposed an approach, scMappR (39), to identify cell type-specific pathways by combining bulk RNA-seq deconvolution with ranked ORA. This method is limited to analyze differentially expressed genes from bulk RNA-seq. Therefore, there is a pressing need for a general tool that allows for convenient cell type-specific gene functional analysis.

The behavior of a cell is determined by the complex interplay of its constituent pathways. Through inter-pathway crosstalk, the gene activity can propagate from one part of the network to others, and the coordination of the entire network determines the development and maintenance of cellular phenotypes. To understand collaboration of genes and pathways in cells, it is necessary to simulate the propagation of signals throughout the entire network, capturing both intra-pathway gene interactions and inter-pathway crosstalk. Existing databases, such as Gene Ontology (40,41) (GO), provide pathway details and their connections, with each of the three biological domains (Biological Process, Molecular Function, and Cellular Component) organizing the directed acyclic graph that resembles the hierarchical organization within a cell. Recent advances in deep learning algorithms have led to the development of novel methods for modeling the hierarchical structure of subsystems (42). In particular, ‘visible’ neural networks (VNNs) have shown promise in learning the signaling propagation within a cell, enabling the dissection of underlying pathways related to specific cellular phenotypes due to its ‘visible’ nature (43,44).

In this study, we developed CellGO, a VNN-based tool for cell type-specific pathway analysis. CellGO integrates the single-cell RNA expression data and the VNN model that emulates the hierarchy of GO terms to capture cell type-specific signatures, intra-pathway gene connections, and inter-pathway crosstalk. We demonstrated the effectiveness of CellGO in identifying cell type-specific active pathways across multiple experimental datasets (14-18) and extended its applicability to a vast collection of 71 publicly available single-cell datasets of human and mouse brains (45-73). CellGO is accessible as a python package cellgopy (https://github.com/FduZhuLab/cellgopy) and through a user-friendly web interface (http://www.cellgo.world).

## 2 Materials and Methods

### Gene Ontology

The Gene Ontology (40,41) depicts a tree-like hierarchical graph where each GO term is a node and edges represent parent-child relations between GO terms. We downloaded the ontology file ‘go-basic.obo’ from https://doi.org/10.5281/zenodo.6799722. This file is the basic version of the ontology, which guarantees the GO graph to be acyclic. We also downloaded the human- and mouse-specific GO annotation files (‘goa_human.gaf.gz’ and ‘goa_human_isoform.gaf.gz’ for human; ‘mgi.gaf.gz’ for mouse) from https://doi.org/10.5281/zenodo.6799722 to species-specifically assign gene products to GO terms. The ontology and annotation data have the release version 2022-7-1 (DOI: 10.5281/zenodo.6799722).

### The CellGO framework

CellGO consists of two phases: (i) The modeling phase includes steps 1-2, developed for scoring cell type-specific activities of gene-term pairs to enable cell type-specific pathway analysis for a single gene, and (ii) The analysis phase includes steps 3-6, developed for comprehensive cell type-specific pathway analysis that works for a gene set. More descriptions of each step are as follows.

#### Step 1: Processing of ontologies and extraction of subtrees

We used the python package goatools (74) v1.0.15 to import the GO graph into the workspace to make it operable and assigned genes to GO terms based on species-specific GO annotations. Then we pruned the species-specifically annotated GO graph according to the following steps: (i) A GO term with less than six annotated genes was deleted, and all genes within it were passed to its first-order parental terms. This procedure was performed from bottom to up to ensure a term is deleted only when it has no child terms. (ii) We deleted the Cellular Component (CC) branch due to its small number of terms. Only the Biological Process (BP) and Molecular Function (MF) branches were retained. After this step, the human- and mouse-specifically annotated GO graph retained 9,870 and 9,979 terms, respectively. (iii) For each non-leaf term (a term with at least one child term), we removed annotated genes within it which simultaneously be annotated in its child terms. (iv) For each non-leaf term, we removed all genes within it if more than 100 genes were annotated. This step avoids the excessive influence of a single term on the global graph. After the above processing of ontologies, we next extracted subtrees from the pruned GO graph. Concretely, for each leaf term with no more than 100 genes, we exported it and all of its parental terms, the parent-child relations between these terms, and the annotated genes of each term as a single file with a specified format. A total of 4,168 and 4,230 subtrees were extracted from the human- and mouse-specifically pruned GO graph, respectively. This part has been coded into a function to facilitate processing other ontology files.

#### Step 2: Scoring cell type-specific activities of gene-term pairs

The VNN was initially inspired by Dcell (43), whose neural network architecture matched the hierarchical structure of the GO graph. Given a subtree including interrelations between terms and annotated genes within each term, CellGO constructs and trains a VNN whose architecture mirrors the inter-pathway parent-child relations and intra-pathway gene annotations. We denote our input training dataset as *D* = {(*X*_1_, *y*_1_), (*X*_1_, *y*_1_),…, (*X*_*N*_, *y*_*N*_)}, where *N* is the number of samples (or cells). For each sample *i, X*_*i*_∈*R*^*G*^ denotes the gene expression, represented as a vector of continuous values on *G* genes, and *y*_*i*_∈*R*^*C*^ denotes the cell identity, represented as a one-hot vector of binary values on *C* classes (or cell types). *y*_*ic*_ = 1 if sample *i* is annotated to class *c*, otherwise *y*_*ic*_ = 0. If an annotated gene does not exist in the single-cell expression matrix, we pad the expression of this gene with zero. The multidimensional state of each term (or pathway) *p*, denoted by the output vector 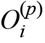, is defined by a nonlinear function of the contribution of its annotated genes *g*^(*p*)^ (if present) and the states of all of its first-order children *child*^(*p*)^, concatenated in the input vector 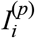:

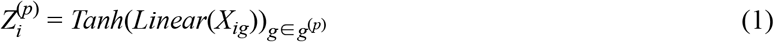

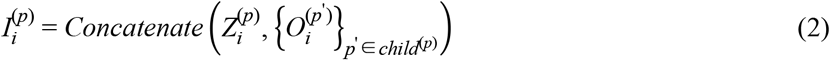

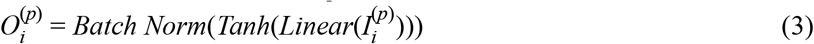

Here, 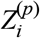 is the contribution vector of annotated genes of *p* with length 10. *Concatenate* returns a single vector that is the concatenation of all input vectors. Let 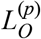 denote the length of 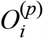, representing the number of values in the state of *p* and determined by

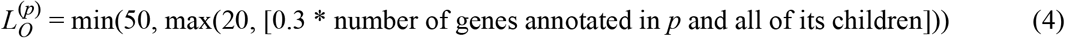

In equation (3), the parameters of *Linear* give the ability to capture the complex biological signals of *p. Tanh* and *Batch Norm* reduce the impact of extreme values and scale differences of weights to accelerate the training speed. We performed the training process by minimizing the objective function:

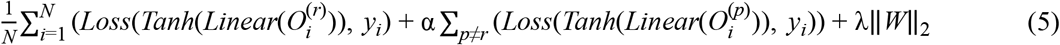

Here, *Loss* is the cross-entropy loss function, and *r* is the top-level term of the subtree. The top-level term can only be one of ‘biological_process’ (GO:0008150) and ‘molecular_function’ (GO:0003674). *Linear* in equation (5) denotes the linear function transforming the multi-dimensional vector 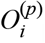 into a vector of continuous values with length *C*. Every term is optimized to predict the cell identities by itself. The parameter α (=0.3) balances these two contributions. λ (=0.001) is an *l*_2_ norm regularization factor. We optimized the objective function using ADAM, and the batch size was selected as one-third of the sample number of the class with the minimum number. We trained each subtree-matched VNN separately using 15 epochs, given its strong learning ability and fast convergence, which also reduced overfitting. We downsampled classes with over 10,000 samples and oversampled classes with less than 500 samples at each epoch (Supplementary Table S1).

After achieving a well-trained VNN corresponding to a subtree *v*, CellGO scores the cell type-specific activities of gene-term pairs within it. Let *gp*∈*GP*^(*v*)^ denote all gene-term paired annotation relations within *v*, and the raw score of a gene-term pair *gp* in class *c* is determined by

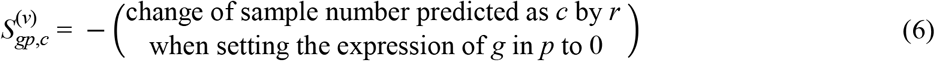

Intuitively, *r* is the parent of all terms in the subtree except itself, which may better respond to the influence of the perturbation of *g* in *p* on the whole subtree. When the number of *c* is decreased then this score is positive and vice versa. We noticed that the impact of the perturbation was transmitted upward along the network architecture to *r*, which simulated the transmission process of biological signals within the hierarchical functional structure of a cell. We next eliminated the class-specific background bias by

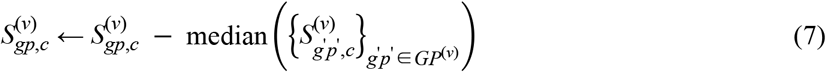

This step allows these scores to reflect relative activity differences between gene-term pairs, and removes the internal bais of the VNN which may cause every score is extreme in a certain class. We next corrected these scores according to the classification accuracy of *r*:

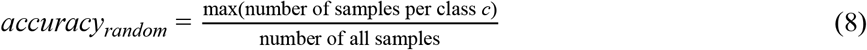

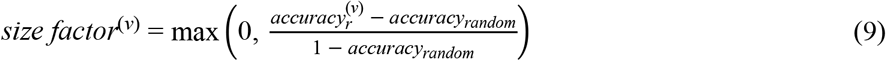

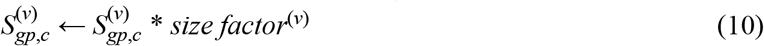

Here, *size factor*^(*v*)^ measures the predictive ability of *r. size factor*^(*v*)^ = 1 if the classification accuracy of *r* equals 1, and *size factor*^(*v*)^ = 0 if the accuracy of *r* is less than the accuracy of classifying all samples to the class with the maximum number. We performed this step because if genes annotated in *v* do not have any cell type-specific signatures, the VNN at this time is not predictable, hence these scores should be rejected. To reduce the time-consuming of the modeling phase, the classification accuracy of *r* is replaced by the classification accuracy for the training dataset rather than the accuracy based on five-fold cross-validation. Let *V* denote the collection of every subtree. Since a *gp* may appear in multiple subtrees, we next computed gene-term scores without consideration of *v*:

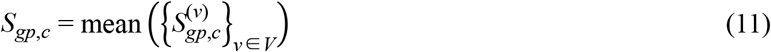

Let *GP* denote the collection of every gene-term pair. We further balanced the power of positive and negative scores by

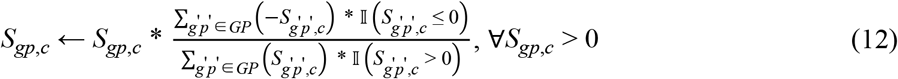

Lastly, we normalized these scores to the interval − 1, 1 by

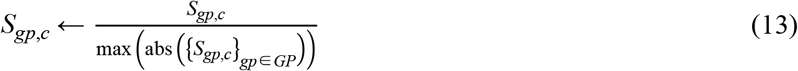

So far, we have completed the estimation of cell type-specific gene-term scores. To facilitate storage and transfer, we exported them as a CSV file. We further defined the *P*-value of a cell type-specific gene-term score by

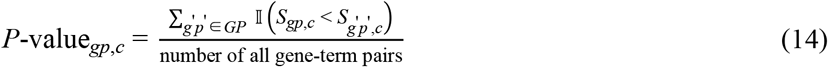

We ran the above pipeline on 71 datasets (Supplementary Table S1) separately to obtain 71 CSV files for downstream analysis. Dataset 1, which includes 25,639 cells and six cell types, spent about 35 hours using one NVIDIA GeForce RTX 2080 Ti GPU.

#### Step 3: Assigning cell type-specific P-values to pathways given a gene list

Given a set of genes denoted by *g*∈*G*^(*u*)^. Let *GP*^(*p*)^ denote the collection of all gene-term pairs which involve term *p* or any one of its child terms, and let *G*^(*p*)^ denote all annotated genes of term *p* and all of its child terms. CellGO next computed cell type-specific pathway active scores (ctPASs) through summing gene-pathway paired active scores (GPPASs):

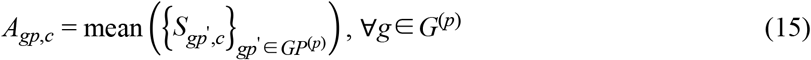

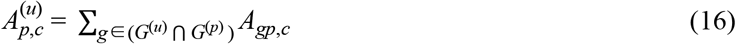

Here, *A*_*gp,c*_ denotes the cell type-specific GPPAS of gene *g* in pathway *p*, if *p* or its children contain *g.* 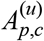 denotes the ctPAS of *p* by summarizing GPPASs of genes in both *G*^(*u*)^ and *G*^(*p*)^. The most active gene in a cell type is defined as the gene with the largest cell type-specific GPPAS. We noticed that a higher GPPAS indicating a more positive and larger contribution to pathway activation. We next generated approximate null distribution of ctPASs by randomly sampling *B* (default 100,000) gene sets 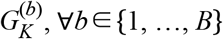, with *K* genes in each:

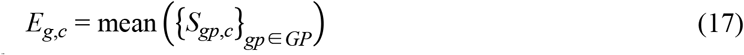

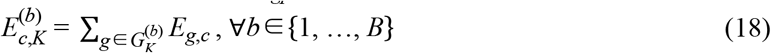

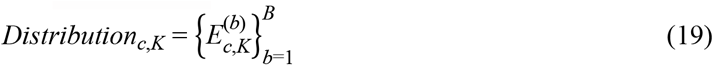

Here, *E*_*g,c*_ denotes the cell type-specific gene active score of gene *g* without consideration of *p*, a universal approximation of *A*_*gp,c*_ in equation (15). 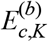 denotes the cell type-specific active scores without consideration of *p. Distribution*_*c,K*_ denotes the null distribution of ctPASs with consideration of gene number *K*. Let 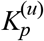 denote the number of genes contained by both *G*^(*u*)^ and *G*^(*p*)^, we assigned the *P*-value to 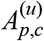 by

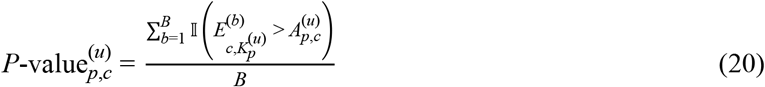

In order to make 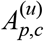 comparable between different pathways and cell types, we calculated the z-scored ctPAS as the output value of the ctPAS by

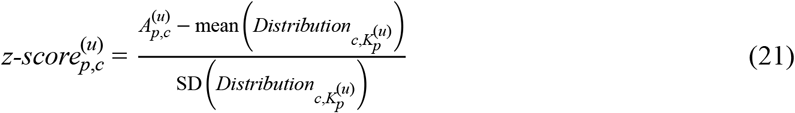

#### Step 4: Analysis of cell-type level activation

We used the python package scipy (75) v1.5.4 to perform Kolmogorov-Smirnov (K-S) test to evaluate the cell type-level significance on a given gene list:

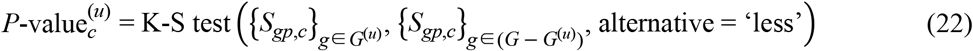

#### Step 5: Cell type-specific pathway network analysis

We screened cell type-specific active pathways by 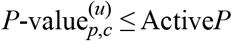 (default 0.01) and took them as seed nodes of random walk with restart (RWR) to explore high-affinity nodes in the pruned GO graph. Since BP and MF are two non-interconnected domains, we implemented RWR on the two domains separately. Let 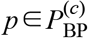 denote active pathways of cell type *c* in BP, we then went on 25 travels starting from each of them, and each travel included ten steps. The first step of each travel was at the starting node. Each of the other nine steps had a 10% probability of returning to the starting and a 90% probability of randomly entering one neighbor (one parent or one child) of the current node. We then selected high-affinity nodes according to the times each node be traveled. The number of high-affinity nodes was defined as [1.1 * number of seed nodes]. It should be noted that high-affinity nodes allow the inclusion of seed nodes.

Let 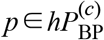 denote high-affinity nodes (or pathways) of cell type *c* in BP, which composed the network of cell type-specific enriched pathways. We then divided them to determine pathway communities. Specifically, we used the python package graph-tool (DOI: 10.6084/m9.figshare.1164194.v14) v2.43 to partition these nodes according to the maximization of Newman’s modularity (76). The graph-tool adopted an agglomerative heuristic to fit the stochastic block model to achieve the community partition. We generated 100 Monte Carlo (MC) partitions and aligned them with a common community labelling. We controlled the lower limit of the number of communities as [0.05 * number of seed nodes]. Let 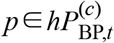 denote pathways of community *t* of cell type *c* in BP, the community-level significance is determined by

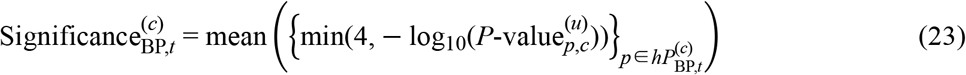

Here, equation (23) denotes the significance of community *t* of cell type *c* in BP. We defined communities with significance greater than − log_10_(Active*P*) (default 2) as active (or significant) communities. We further filtered out communities with a low number of pathways by the average number of pathways contained by significant communities. Prioritized top communities were selected based on significance rankings. In this study, we manually summarize the topic of a community to maximally cover the semantics of pathways within the community. It should be noted that all results of this part reported in this paper were based on BP because MF contained a low number of pathways. For the differentially expressed gene (DEG) list generated from the Conrow-Graham et al. *Adnp* KO dataset (17), we set Active*P* to 0.001 because there were more than 500 original active pathways in microglia.

#### Step 6: ORA-based pathway analysis in CellGO

CellGO included the function of conventional pathway enrichment analysis similar to gProfileR (25-27). Given a set of genes denoted by *g*∈*G*^(*u*)^, and let *G*^(*p*)^ denote the collection of all annotated genes of a term *p* and all of its child terms. We next used the python package scipy (75) v1.5.4 to perform Fisher’s exact test to evaluate the pathway-level significance on a given gene list:

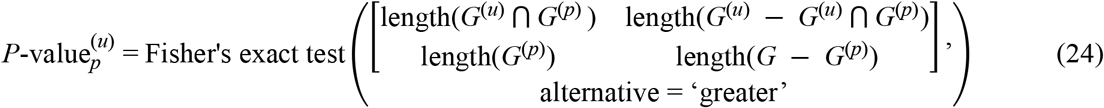

These results were output simultaneously with cell type-specific *P*-values.

### Embedding a set of pathways to the semantic space

Given a set of pathways denoted by *p*∈*P*^(*u*)^, where *P* represents the collection of every pathway *p* in the pruned GO graph. The probability of a pathway is defined as

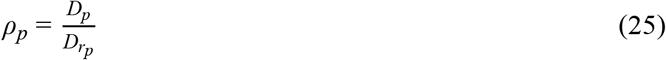

Here, *D*_*p*_ denotes the number of *p* and all of its children. *r*_*p*_ denotes the top-level pathway of *p*. The top-level pathway can only be one of ‘biological_process’ (GO:0008150) and ‘molecular_function’ (GO:0003674). Let *ρ*_min_(*p*_1_, *p*_2_) denote the minimum probability of the common ancestor pathways of *p*_1_ and *p*_2_, we next used the Lin method (77) to determine the similarity between the two pathways:

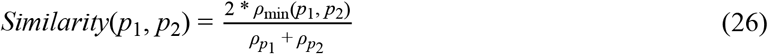

We computed the similarity for all pathway-pathway pairs in *P*^(*u*)^, which composed the similarity matrix ℝ^*U*×*U*^, where *U* is the number of pathways in *P*^(*u*)^. We next used the umap function from the R package umap (78) v0.2.9.0 with default parameters to reduce the length of the second dimension of the similarity matrix to two. Since BP and MF are two non-interconnected domains, we only mapped the BP pathways to the two-dimensional semantic space.

### Identification of predefined cell type-specific pathways

We collected marker genes of five cell types from the McKenzie et al. study (79). These five cell types included neurons, oligodendrocytes, oligodendrocyte precursor cells, astrocytes, and microglia (Supplementary Table S2). The top 50 marker genes were selected for each cell type based on the reported rankings. We then took the marker genes of each cell type as input to perform conventional pathway enrichment analysis using CellGO. Predefined cell type-specific pathways were selected for each cell type according to the top 50 pathways ranked by *P*-values (Supplementary Table S3).

### Collection and processing of single-cell RNA-seq datasets

Human and mouse brain scRNA-seq or snRNA-seq datasets (45-73) were collected from publicly available resources, including GEO (https://www.ncbi.nlm.nih.gov/geo/), UCSC Cell Browser (https://cells.ucsc.edu/), Broad Single Cell Portal (https://singlecell.broadinstitute.org/), Human Cell Atlas (https://data.humancellatlas.org/), Allen Brain Atlas (http://celltypes.brain-map.org/download), DropViz (http://dropviz.org/), and Synapse (https://www.synapse.org/). We only selected datasets in which both the processed expression data and pre-assigned cell labels were available. Processed expression data could be read count and UMI count. It could also be CPM, RPKM, TPM, and log-transformation of these forms. We did not perform any pre-analysis on FASTQ files.

A total of 71 datasets were curated and processed separately to make up the proposed database according to the following steps: (i) When the obtained value was read count or UMI count, it will be normalized by the NormalizeData function from the R package Seurat (80) v4.3.0 with default parameters to allow correction for the total number of reads per cell and log-transformation. Other values, such as CPM, RPKM, and TPM, were log2-transformed with pseudo-count one unless the expression data was already log-transformed. (ii) Quality control (QC) of cells was performed as described in the original study unless the obtained dataset was already QC-ed. (iii) Cells with uninformative cell type labels (e.g., ‘unclassified’) defined as outliers in the original study were excluded. We also removed unimportant and low-quantity cell types in the brain, such as T cells, endothelial cells, pericytes, and vascular leptomeningeal cells. We only retained cells of the control group. (iv) We manually aligned cell type labels annotated in the original study to the unified terminology to ensure consistency between datasets. 61 out of the 71 datasets were re-annotated at major-type-level, including excitatory neurons, inhibitory neurons, radial glias, intermediate progenitor cells, etc. Other ten datasets, which included only one major cell type, were re-annotated at subtype-level, such as excitatory neuron subtypes, including L4 IT, L5/6 IT, L6b, etc. (v) Re-annotated cell types with the number of cells less than 100 were removed. For each dataset, we exported the re-processed expression matrix and the cell identity list as two CSV files and took them as input to the modeling phase of CellGO. Details about each dataset, such as references, original download URLs, species, anatomical areas, and included cell types, are reported in the Supplementary Table S1.

### Collection of experimental DEG lists

We first considered five experimental datasets, including the bulk RNA sequencing data for both the experimental and control groups (Supplementary Table S4). We next screened the top 100 differentially expressed genes based on *P*-value rankings from the supplementary data of corresponding references. Differentially expressed genes were achieved by differential gene expression analysis between the experimental and control groups. For the Conrow-Graham et al. *Adnp* KO dataset (17), the supplementary data only reported up-regulated genes in the KO group.

### Comparison with other tools

We compared our method with two existing methods: gProfileR (25-27) and scMappR (39). We applied these two methods to DEG lists from Smith et al. (14), Runge et al. (15), and Wang et al. (16). gProfileR takes a set of genes as input and returns *P*-values of pathways. We used the gprofiler function from the R package gProfileR v0.7.0 with default parameters to analyze the collected DEG lists and obtained FDR-corrected *P*-values. scMappR requires not only a DEG list but also the *P*-values and log-transformed fold changes of DEGs, the bulk expression matrix, and the single-cell gene signature matrix as input, then scMappR returns cell type-specific *P*-values of pathways. The single-cell gene signature matrix was generated based on dataset 47 using the seurat_to_generes and generes_to_heatmap functions from the R package scMappR v1.0.9 with default parameters. The adopted type of the single-cell gene signature matrix was ‘OR’ (odds ratios). The *P*-values and log-transformed fold changes matching the collected DEG lists were obtained from the supplementary data of corresponding references (Supplementary Table S4). The bulk expression matrices were collected from GEO (Data Availability). We next took them to the scMappR_and_pathway_analysis function with default parameters and obtained cell type-specific FDR-corrected *P*-values. It should be noted that we only retained pathways with the domain BP or MF.

## 3 Results

### Design of CellGO

CellGO is a tool designed to identify cell type-specific active pathways impacted by a single gene or a gene set, operating in two phases: a modeling phase and an analysis phase (Figure 1).

**Figure 1.**
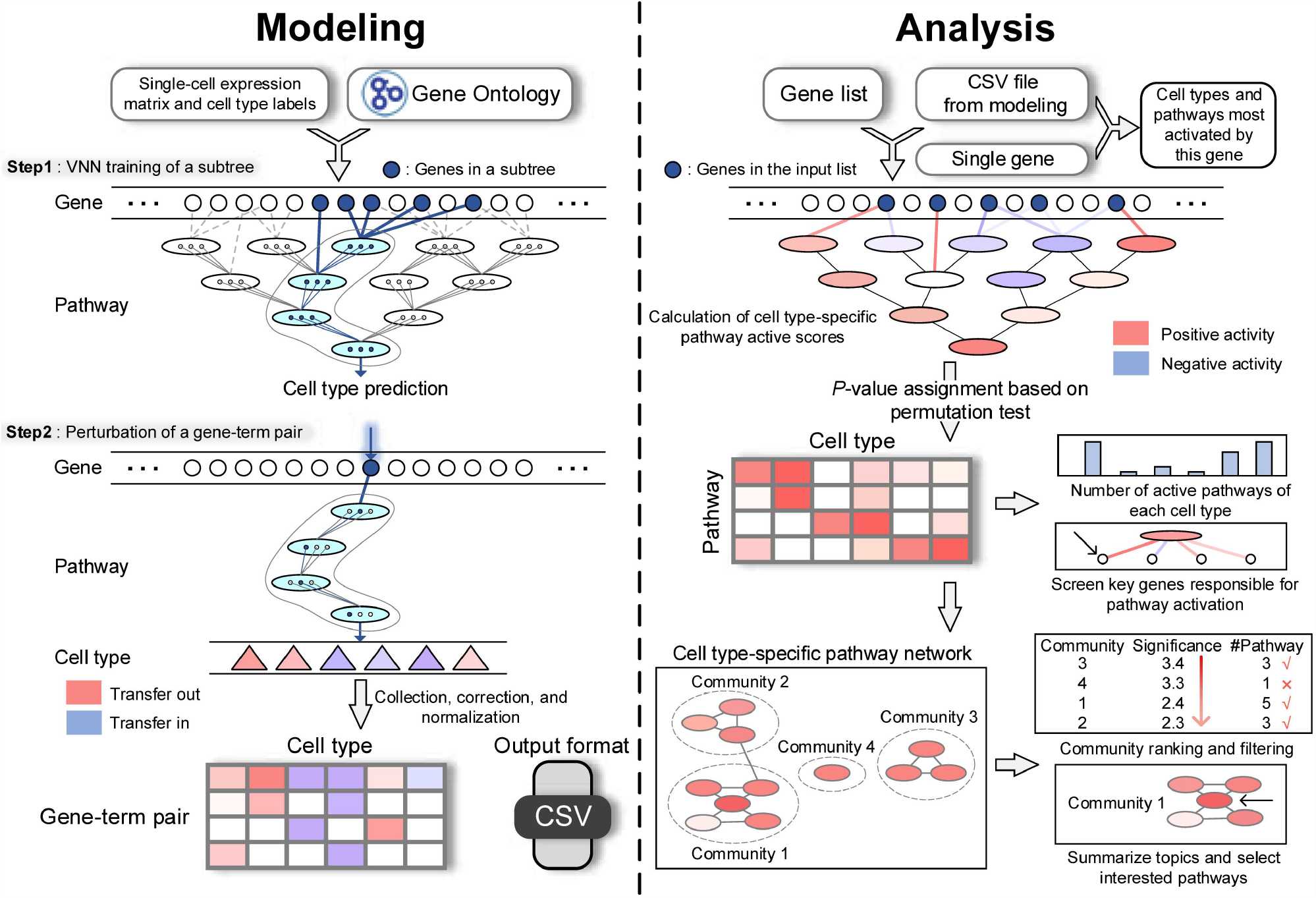
CellGO workflow. CellGO predicts cell type-specific active pathways of a single gene or a gene set through a modeling phase (left) and an analysis phase (right). In the modeling phase, CellGO utilizes the single-cell gene expression matrix to learn cell type labels by training VNNs based on subtrees starting from each leaf term (Step 1). It then scores the cell type-specific activity of each gene-term pair as GTSs by quantifying changes in cell type labels when simulating gene perturbation. Cell type-specific GTSs are reported as a CSV file for subsequent analysis. In the analysis phase, CellGO employs a novel pathway activation measuring system to infer cell type-specific active pathways in the single-gene mode, and a four-level hierarchy of biological interpretation in the gene-set mode.

During the modeling phase, the cell type-specific signaling transduction process within a cell is simulated using single-cell gene expression data to predict cell type labels through the training of a VNN with a structure that mirrors the hierarchy of GO terms. The VNN used in CellGO is a non-fully connected feedforward neural network, where neurons of a GO term receive inputs solely from neurons of its direct child terms and the genes present in this term but not in its child terms (43). To capture the multi-function of each term, CellGO represents each term with multiple neurons ranging from 20 to 50. CellGO constructs a global VNN using all GO terms. Additionally, CellGO constructs a subtree for each leaf term (a term without any child terms), consisting of the leaf term and all its parental terms that are directly or indirectly connected to it, and learns cell type labels using this subtree. The gene-term score (GTS), which represents the paired active score between a gene and the term it directly connects to, is determined by simulating the gene expression to zero and measuring the resulting alteration of cell identities (Materials and Methods). The modeling phase outputs a CSV file containing cell type-specific GTSs of all gene-term pairs for the subsequent analysis phase.

The analysis phase takes the CSV file generated from the previous modeling phase, along with either a single gene or a gene set for query as input. CellGO is capable of predicting the cell types and cell type-specific pathways significantly activated by a single gene by calculating empirical *P*-values for GTSs (Materials and Methods). For a gene set, gene-pathway paired active scores (GPPASs) are then calculated based on GTSs. GTSs are assigned as GPPASs for directly connected gene-pathway pairs. When a gene and a pathway is indirectly connected, the GPPAS is computed by averaging the GTSs of the pathway’s all direct and indirect children that are directly connected to the gene. The cell type-specific pathway active score (ctPAS) is obtained by summing the GPPASs of all genes in both the pathway and the set. To identify cell type-specific active pathways, CellGO compares ctPASs of the query set with the empirical distribution of ctPASs calculated by randomly sampling the same number of genes (Materials and Methods).

The analysis phase offers several modules for further mechanistic interpretation of the query gene set. Firstly, it can select cell type-specific active pathways based on empirical *P*-values and z-scored ctPASs. Secondly, it can identify the cell type(s) most activated by the enquiry gene set by either ranking the number of active pathways or by comparing the distribution of GTSs with the background utilizing Kolmogorov-Smirnov (K-S) test across cell types. Thirdly, CellGO can construct the network of cell type-specific active pathways and report top communities enriched with active pathways, by incorporating the random walk with restart algorithm (81) and the community partition algorithm. In this study, we manually summarize the topic of a community to maximally cover the semantics of pathways within the community. Lastly, CellGO allows for the selection of the most active gene (MAG) for each active pathway based on GPPASs.

### A comprehensive database to demonstrate the application of CellGO

To demonstrate the applicability of CellGO, we implemented our model on 71 single-cell datasets (45-73), comprising over three million cells from 47 prenatal and postnatal brain regions and 34 cell types (Figure 2A and Supplementary Table S1). We provide a web interface for interactive querying of cell type-specific pathway analysis and visualization of results, and all CSV files generated from the modeling phase are available for download to enable local analysis using the python package cellgopy.

**Figure 2.**
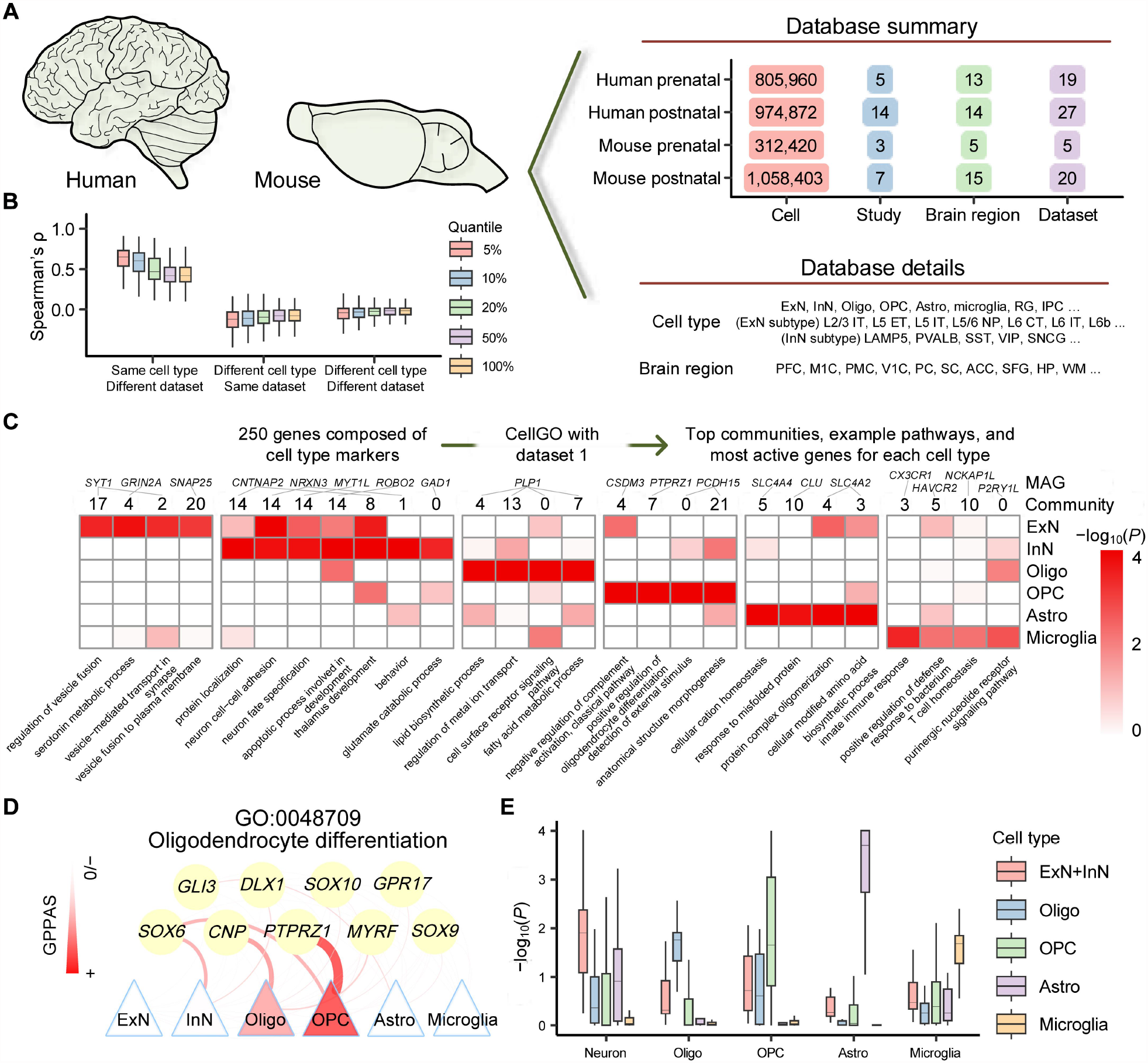
Application of CellGO to capture cell type-specific signatures with built-in datasets. (**A**) Overview of brain cell datasets inbuilt in CellGO. The numbers of cells, studies, brain regions, and datasets are shown in the upper right. Representative cell types and brain regions are listed in the lower right. The full names of cell types are listed in the Supplementary Table S1. (**B**) Boxplot showing pairwise Spearman’s rank correlation coefficients across GTSs of the same cell type from different datasets (left), GTSs of different cell types from the same dataset (middle), and GTSs of different cell types from different datasets (right), calculated using 27 human postnatal datasets. Spearman’s correlations calculated using different quantiles of gene-term pairs were displayed. For example, 5% indicates that the correlations were calculated on gene-term pairs with GTSs ranked in the top 5% in either group. The horizontal middle line of each boxplot denotes the median. The bounds of the box represent 0.25 quantile (*Q*_1_) and 0.75 quantile (*Q*_3_), respectively. The upper whisker is the minimum of the maximum value and *Q*_3_ + 1.5\**IQR*, where *IQR* = *Q*_3_ − *Q*_1_. The lower whisker is the maximum of the minimum value and *Q*_1_ − 1.5\**IQR*. (**C**) Top four communities, example pathways in these communities, and MAGs of these pathways for each cell type identified by CellGO testing 250 cell type markers using dataset 1. Heatmap colors denote the cell type-specific log-transformed *P*-values of example pathways in the top four communities per cell type. The community index and the MAG of each pathway are shown above the heatmap. (**D**) Activation map showcasing genes in the oligodendrocyte differentiation pathway across different cell types. The thickness and colors of lines denote GPPASs (Materials and Methods). (**E**) Box plot showing the distribution of cell type-specific log-transformed *P*-values from CellGO for predefined cell type-specific pathways. ‘ExN+InN’ in the legend denotes the maximum value of the minus log-transformed *P*-values of ExN and InN.

### Reproducibility and robustness

We first tested the prediction accuracy of CellGO on cell type classification based on dataset 1 that includes 25,639 cells and six cell types of the adult neocortex (Supplementary Figure S1). The global VNN, which matched the entire GO graph, achieved a cell type classification accuracy of 99.6% according to the results repeating five-fold cross-validation for five times. Additionally, we trained models based on 4,168 VNNs matched with subtrees that included a median of ten pathways and 99 genes. These subtree-based models achieved a median accuracy of 78.7% in cell type classification.

As a stochastic learning algorithm, a feedforward neural network produces different optimized parameters each time it is run. To assess the reproducibility of CellGO’s neural networks, we performed the modeling phase four times on dataset 1. We found high correlations of GTSs between replicates of the same cell type (Supplementary Figure S2). Specifically, considering gene-term pairs ranking in the top 5% in either of the two replicates, the median Spearman’s ρ achieved 0.904, and 0.785 for all gene-term pairs. Conversely, we observed very low correlations of GTSs between different cell types within (median Spearman’s ρ = −0.0537) and between (median Spearman’s ρ = −0.0391) replicates, indicating distinct gene-term activation patterns among cell types. Similar patterns were observed in human postnatal datasets (Figure 2B), albeit with lower correlations for both top 5% gene-term pairs (median Spearman’s ρ = 0.665) and all gene-term pairs (median Spearman’s ρ = 0.433) between different datasets of the same cell type. These lower correlations could be partially attributed to differences in cell types across brain regions and batch effects between datasets. Notably, the correlations among different datasets of the same cell type are considerably greater than those among different cell types within the same dataset (Figure 2B). Overall, these results indicate that CellGO generates replicable and robust GTSs for downstream analysis.

### Deconvolution of cell type-specific signatures

To assess the ability of CellGO to accurately identify cell type-specific pathways, we assembled a query gene set consisting of the top 50 marker genes for each cell type from an independent study (79) (Supplementary Table S2). The significant pathways identified by Fisher’s exact test, using each cell type’s marker genes independently, were considered as ground truth of predefined cell type-specific pathways (Supplementary Table S3; Materials and Methods). Our analysis using the assembled gene set identified top communities enriched with active pathways with distinct semantics in different cell types (Figure 2C and Supplementary Table S5). In addition, CellGO assigned more significant *P*-values for predefined cell type-specific pathways in their corresponding cell types (Figure 2E). All these results were further corroborated by another dataset (Supplementary Figure S3 and Supplementary Table S6). Overall, these findings suggest that CellGO is effective in identifying cell type-specific pathways and extracting cell type-specific features from a gene set.

Moreover, CellGO is able to identify MAGs in cell type-specific active pathways (Materials and Methods). For example, in excitatory neurons (ExN), the gene *SYT1* that encodes a membrane protein involved in triggering neurotransmitter release at the synapse, was identified as the MAG in pathways, such as regulation of vesicle fusion and vesicle-mediated transport in synapse, belonging to top communities (Figure 2C). In oligodendrocytes, we indicated *PLP1*, a gene encoding a transmembrane proteolipid protein that is the predominant component of myelin, as the MAG in lipid biosynthetic process and regulation of metal ion transport. Notably, CellGO determines MAGs of the same pathway in a cell type-specific manner. For instance, CellGO identified *PTPRZ1* as the MAG of oligodendrocyte differentiation in oligodendrocyte precursor cells (OPC) (Figure 2D), consistent with its reported role in maintaining OPC in an undifferentiated state (82). In contrast, *CNP*, a marker of late-phase OPC, was inferred to be the MAG of the same pathway in oligodendrocytes, aligning with its known function in the retention of the myelin sheath (83).

Together, the above results demonstrate that CellGO is an effective tool for identifying cell type-specific active pathways from a gene set and key genes that most activate these pathways. By combining pathway-level and gene-level analysis, CellGO can provide insights into the multicellular mechanisms as well as facilitate the identification of driving genes.

### Functional inference of gene knockouts and comparative experiments

The performance of CellGO on cell type-specific pathway analysis was further assessed using gene knockout (KO) datasets (14-15,17-18) (Supplementary Table S4). First, we applied CellGO to examine *Atp1a2*, a gene encoding α2-Na/K ATPase that is responsible for ATP binding activity and maintenance of the ion electrochemical gradients across the membrane organization, using single-gene analysis based on the model trained on a mouse postnatal primary visual cortex dataset (dataset 47). Our analysis revealed that astrocytes were the most affected cell type by *Atp1a2* perturbation, displaying the most significant *P*-values of GTSs compared to other cell types (Figure 3A). Pathways with the highest GTSs in astrocytes were related to ion homeostasis and stimulus responses (Figure 3B and Supplementary Table S7), matching previous experimental studies associating *Atp1a2* with astrocyte-mediated K and Ca ion transport and LPS-induced infammation (84-86). In contrast, pathways with the lowest GTSs in astrocytes were associated with neuron-dependent processes, such as learning, behaviors, and neurotransmitter uptake (Supplementary Table S7). As analysis of a single gene largely depends on its annotation in the GO database, we extended our analysis to the top 100 differentially expressed genes between astrocyte-specific *Atp1a2* KO and WT mice to gain more detailed insights (14). Similar to the results from the single-gene mode, CellGO revealed the largest number of active pathways in astrocytes compared to other cell types. Noteworthily, three out of the top five communities of astrocytes represented topics related to metabolic processes (Supplementary Table S8). More specifically, ATP metabolic process, amine biosynthetic process, and primary amino compound metabolic process were discovered in these communities (Figure 3C), with *Atp1a2, Sat1*, a gene encoding an enzyme involved in the polyamine metabolism, and *Cyp2d22*, a gene encoding an enzyme that participates in wide metabolic and biosynthetic processes, as the MAGs in these pathways, respectively. These findings align with a previous study (14) indicating that *Atp1a2* deletion in astrocytes leads to alterations of transcript levels of metabolic enzymes and metabolite levels of serine and glycine, as well as impairments in mitochondrial function, establishing a link between *Atp1a2* and amino acid metabolism in astrocytes within the neocortex.

**Figure 3.**
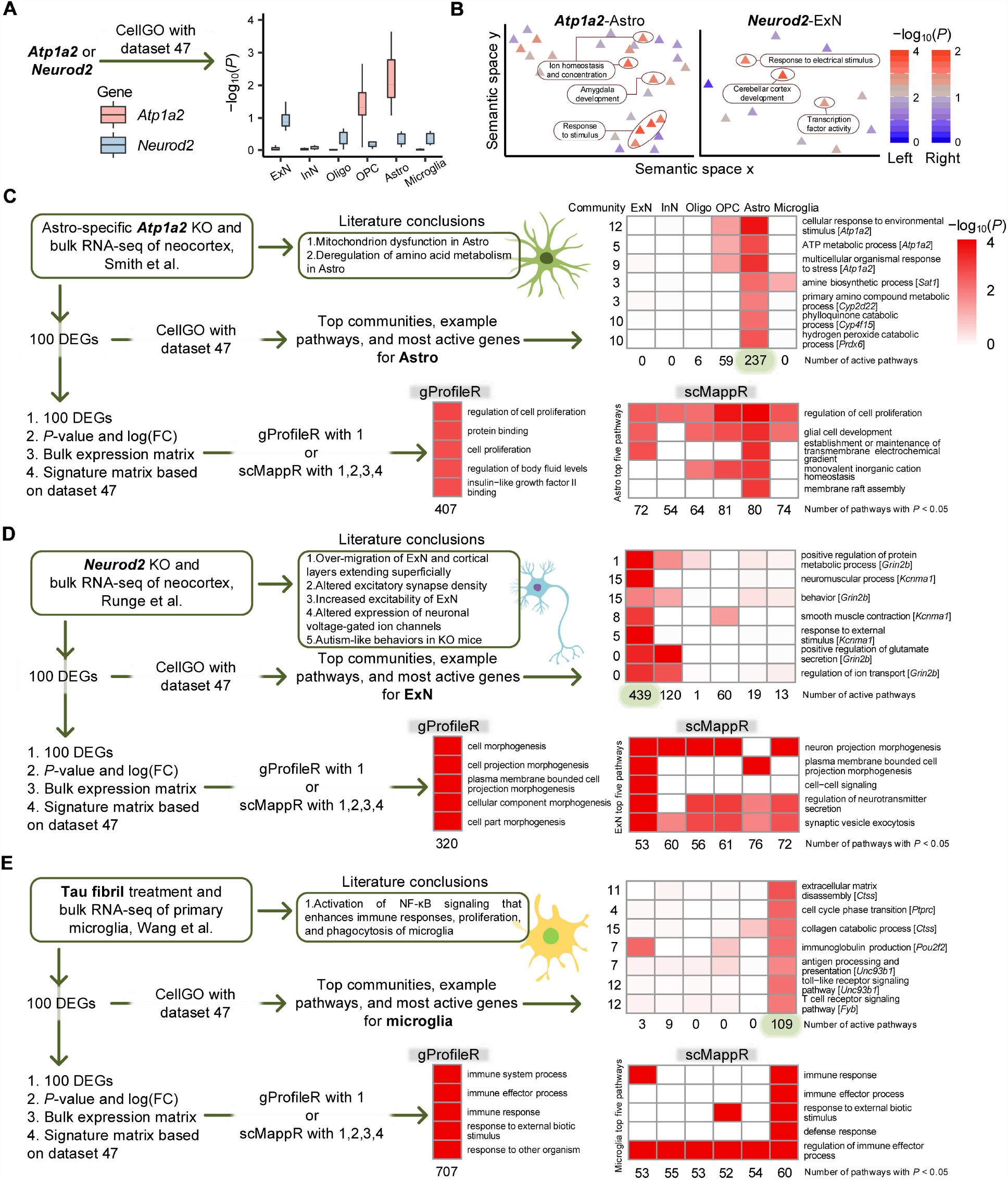
Benchmarking CellGO’s performance on experimental datasets. (**A**) Boxplot illustrating the distribution of cell type-specific log-transformed *P*-values, mimicking *Atp1a2* and *Neurod2* knockout, using single-gene mode based on dataset 47 (Materials and Methods). The boxplot displays the median as the horizontal middle line, the 0.25 quantile (*Q*_1_) and 0.75 quantile (*Q*_3_) as the bounds of the box, and the upper and lower whiskers based on the minimum of the maximum value and *Q*_3_ + 1.5\**IQR*, and the maximum of the minimum value and *Q*_1_ − 1.5\**IQR*, respectively. (**B**) Semantic map of pathways predicted to be perturbed in *Atp1a2* (left) and *Neurod2* (right) knockouts, respectively, using single-gene mode. The colors of triangles denote the log-transformed *P*-values of pathways directly connected to *Atp1a2* in astrocytes (left) or log-transformed *P*-values of pathways directly connected to *Neurod2* in ExN (right). The most significant pathways are elliptically labeled and annotated with their names. The embedding of pathways into the semantic space is described in the Materials and Methods. (**C**,**D**,**E**) Comparison of results obtained from CellGO, gProfileR, and scMappR when analyzing differentially expressed gene lists from (**C**) Smith et al., (**D**) Runge et al., and (**E**) Wang et al.. The main conclusions reported in the original studies are listed as ground truth. Heatmap colors represent log-transformed *P*-values for CellGO and FDR-adjusted *P*-values for gProfileR and scMappR. In CellGO heatmaps, example pathways in the top five communities of the most active cell type are plotted. The number of active pathways (*P*-value < 0.01) per cell type is displayed underneath the heatmap, and the MAG of each pathway is provided next to the pathway name. The gProfileR heatmaps depict the top five significant pathways, with the number of significant pathways (FDR-adjusted *P* < 0.05) shown underneath the heatmap. The scMappR heatmaps present the top five significant pathways in the same cell types as the corresponding CellGO heatmaps, with the number of significant pathways (FDR-adjusted *P* < 0.05) of each cell type labeled underneath the heatmap. FC, fold change. DEG, differentially expressed gene.

For comparison, we also analyzed the top 100 differentially expressed genes using gProfileR (25-27) and scMappR (39). The top enriched pathways identified by gProfileR included regulation of cell proliferation, protein binding, and regulation of body fluid levels (Figure 3C and Supplementary Table S8). Notably, gProfileR discovered 407 enriched pathways in BP and MF domains, a larger number than astrocyte-active pathways identified by CellGO, as gProfileR is not designed to provide cell type information. Pathways revealed by gProfileR included some with semantics that align with the conclusions of experimental studies (14), such as cellular modified amino acid catabolic process and potassium-transporting ATPase activity, although these pathways are ranked below the top 20 in gProfileR results. On the other hand, unlike CellGO, scMappR identified the largest number of significant pathways in OPC (Figure 3C and Supplementary Table S8), and displayed a lower level of cell type specificity in enriched pathways compared to CellGO. Specifically, 84% (357 out of 425) of the pathways enriched in a certain cell type were also enriched in other cell types in scMappR, whereas only 8% (24 out of 302) of active pathways were found to be shared across multiple cell types in CellGO. Moreover, none of the aforementioned experimentally verified pathways were reported as significant in astrocytes by scMappR.

Next, we utilized CellGO to predict the effect of *Neurod2* KO on cell type-specific pathways. Single-gene analysis revealed that *Neurod2* primarily affected ExN (Figure 3A), with most influenced pathways related to electrical stimulus responses, cerebellar cortex development, and transcription factor activity (Figure 3B). Expanding the analysis to the top 100 differentially expressed genes between *Neurod2* KO and WT mice (15), CellGO identified ExN as the cell type with the highest activation that has 439 active pathways (Figure 3D). The top communities in ExN represented topics related to nervous and muscle system processes, secretion, behaviors, stimulus responses, and transport (Supplementary Table S8), and specific pathways within them are shown in the Figure 3D, with *Grin2b*, a gene encoding a protein that acts as an agonist binding site for glutamate, predicted as the MAG of multiple pathways. Other communities significantly enriched with ExN-active pathways possessed various topics associated with cell death, cell differentiation, anatomical structure morphogenesis, ion homeostasis, and membrane potential (Supplementary Figure S4A). These findings are consistent with previously reported functions of *Neurod2* on ExN differentiation and maturation in the neocortex (15,87). KO of *Neurod2* leads to excessive migration of ExN, resulting in alterations of cortical layer sizes and positionings (15). Additionally, *Neurod2* depletion also induces dysregulated expression of ion channels, elevation of ExN action potentials in response to depolarizing current injections, as well as the manifestation of autism-like behaviors (15).

In contrast to the results obtained from CellGO, analysis of the top 100 differentially expressed genes by gProfileR highlighted top enriched pathways related to cell morphogenesis (Figure 3D and Supplementary Table S8). On the other hand, scMappR identified astrocytes as the cell type with the largest number of significant pathways (Figure 3D and Supplementary Table S8). Top enriched pathways in ExN computed by scMappR included neuron projection morphogenesis, plasma membrane bounded cell projection morphogenesis, cell-cell signaling, regulation of neurotransmitter secretion, and synaptic vesicle exocytosis, with four out of these five pathways being significant across multiple cell types.

The consistency between CellGO’s predictions and experimental validations was further substantiated by analyzing additional *Adnp* KO (17) and *Wwox* knockdown (18) datasets (Supplementary Figure S4B-C; Supplementary Text; Supplementary Table S8). Furthermore, we demonstrated CellGO’s implementation in analyzing differentially expressed genes produced from comparative experiments between chemical treatment and normal control. Our analysis of primary microglia treated with tau fibrils (16) revealed that microglia exhibited the highest activation (Figure 3E), with the top communities of microglia representing topics related to immune and metabolic processes, extracellular structure organization, and cell cycle (Supplementary Table S8). These communities included pathways such as extracellular matrix disassembly, cell cycle phase transition, immunoglobulin production, and toll-like receptor signaling pathway (Figure 3E), with *Ctss, Ptprc, Pou2f2*, and *Unc93b1* identified as the MAGs in these pathways, respectively. These findings align with previous experimental studies (16) revealing that tau activates the NF-κB signaling that enhances microglial immune responses, proliferation, and phagocytosis. Both gProfileR and scMappR analyses also identified top enriched pathways related to immune responses (Figure 3E and Supplementary Table S8). While microglia displayed the largest number of significant pathways according to scMappR, every cell type showed enrichment of numerous immune pathways.

Overall, the above results demonstrated that CellGO outperformed the existing tools in identifying functionally relevant cell types and cell type-specific pathways. Moreover, CellGO’s ability to perform single-gene-based analysis and to identify the MAGs in critical pathways from the query gene set provides unique insights into molecular mechanisms, distinguishing it from other pathway analysis tools.

### Cell type-specific pathway analysis of disease risk genes

Large-scale GWASs have been pivotal in identifying risk genes for various diseases (6-13). However, understanding the underlying pathogenic mechanisms remains challenging, despite efforts to infer disease-associated cell types and biological processes through the integration of GWAS summary statistics and single-cell RNA sequencing data (88-90). To address this, we applied CellGO to investigate the cell type-level pathogenesis of Alzheimer’s disease (AD), autism spectrum disorder (ASD), and Parkinson’s diseacse (PD), by analyzing risk genes using datasets from the primarily affected brain regions of these disorders.

CellGO was first applied to analyze 76 risk genes of AD (6) (Supplementary Table S9) using a human entorhinal cortex (EC) dataset (dataset 4), as EC is known to exhibit the earliest histological alterations in AD (91). In accordance with a recent study (92), our analysis identified microglia and oligodendrocytes as the most closely associated cell types in this region, in terms of both the number of active pathways and cell type-level enrichment based on K-S test (Figure 4A; Materials and Methods). The top communities of microglia represented topics related to proteolysis, amyloid-beta formation, lipid and protein metabolic processes, signal transduction, and myeloid cell differentiation (Figure 4B and Supplementary Table S10), consistent with previously reported findings in AD patients where abnormal microglial responses to amyloid-beta are associated with the accumulation of amyloid plaques (93), and altered protein and lipid metabolism in microglia with AD pathology (94-96). In addition, we determined *SORL1* as the MAG of pathways across multiple top communities (Figure 4B), aligning with its known function in amyloid-beta destruction (97). The top communities of oligodendrocytes revealed biological processes distinct from those in microglia (Figure 4C), characterized by topics related to metal ion transport and responses, membrane potential, and gliogenesis. These findings corroborate existing results revealed by single-cell analysis that genes related to ion transmembrane transport and membrane potential were down-regulated in oligodendrocytes of AD patients compared with healthy controls (98). Furthermore, we discovered *APP*, a gene encoding the amyloid precursor protein, as the MAG of pathways in communities associated with metal ion transport, protein metabolic processes, and organic substance responses (Figure 4C). *APP* is known to play an essential role in AD, where accumulation of the amyloid-beta peptides from proteolytic cleavage of the amyloid precursor protein is the key factor in dysregulation of the ion transport process and the development of AD (99,100). Therefore, CellGO can not only uncover cell type-specific functional pathways that contribute to AD but also identify the key genes that drive these pathogenic mechanisms.

**Figure 4.**
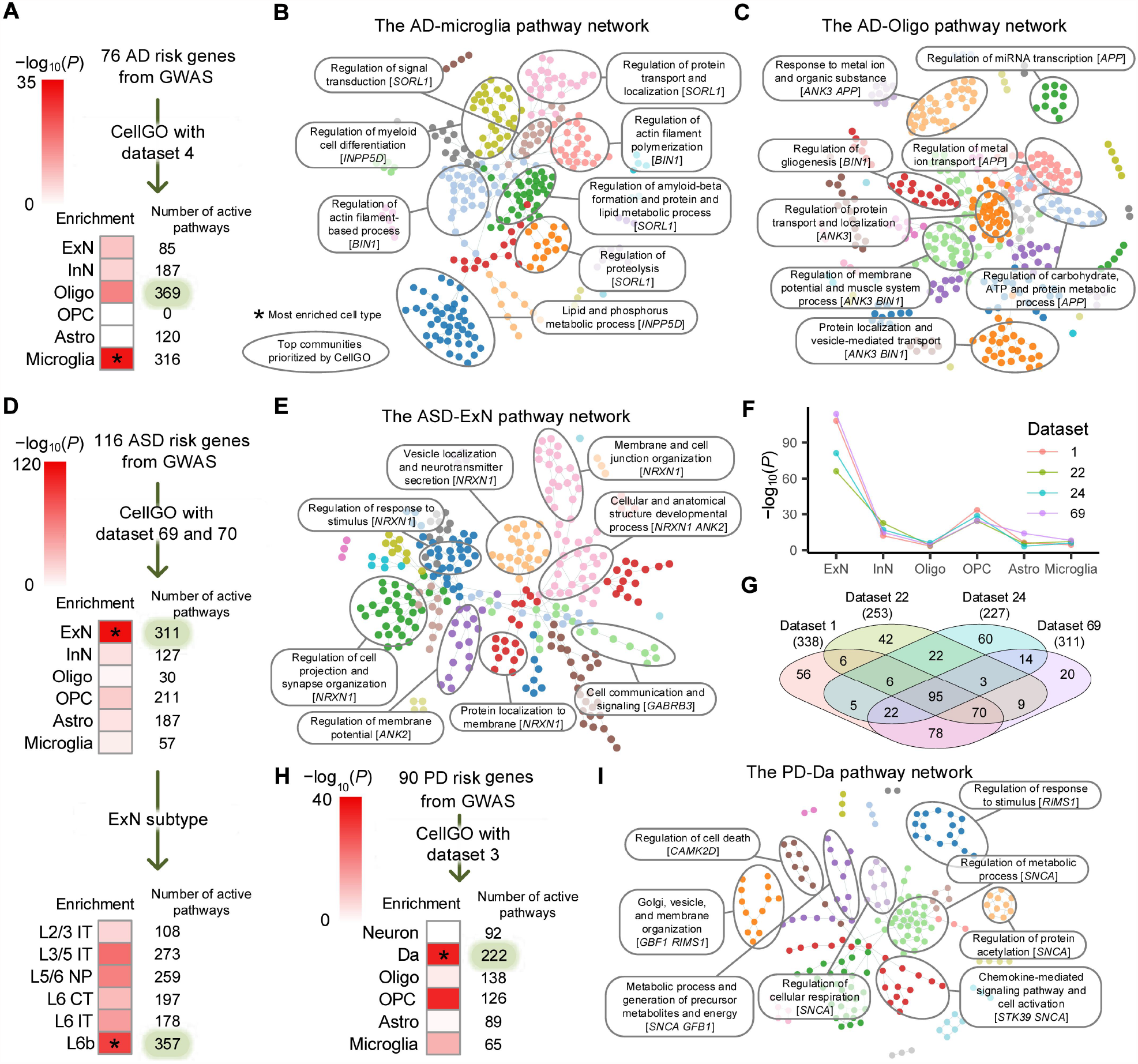
Application of CellGO to disease risk genes. (**A**,**B**,**C**) Results of CellGO analysis performed on 76 AD risk genes using dataset 4. (**A**) The cell type enrichment of AD-active pathways. Heatmap colors denote the log-transformed *P*-values of cell types based on K-S test of GTSs (Materials and Methods). The asterisk denotes the most enriched cell type. The number of active pathways (*P* < 0.01) of each cell type is shown on the right side of the heatmap. (**B**,**C**) The network of AD-active pathways in (**B**) microglia and (**C**) oligodendrocytes. The pathways are colored by communities, with the top eight communities elliptically labeled and annotated with topics. MAGs of pathways with close semantics to the topic of communities are labeled. (**D**,**E**,**F**,**G**) Results of CellGO analysis performed on 116 ASD risk genes. (**D**) The cell-type and ExN subtype enrichment of ASD risk genes based on dataset 69, respectively and 70. (**E**) The network of ASD-active pathways in ExN. (**F**) Cell-type enrichment of pathways activated by ASD risk genes using four different PFC datasets. Log-transformed *P*-values of K-S test based on GTSs were plotted. (**G**) Venn diagram showing the numbers of shared ASD-associated ExN-active pathways between four PFC datasets. The total number of ExN-active pathways per dataset is shown in parentheses. (**H**) The cell type enrichment of pathways activated by 90 PD risk genes using dataset 3. (**I**) The network of PD-associated pathways in Da.

Next, we applied CellGO to study 116 ASD risk genes (7-10) using a prefrontal cortex (PFC) dataset (dataset 69). We found that ExN, particularly deep-layer subtypes, exhibited the highest level of activation (Figure 4D), consistent with previous experimental analysis (101). ExN’s top communities possessed topics associated with synapse and membrane organization, vesicle localization, membrane potential, signaling, and developmental processes (Figure 4E). Similar results were found in the pathway network of the most active ExN subtype, L6b (Supplementary Figure S5). These findings are in line with previous studies that have revealed the impairment of synaptic development and functions in ASD (45,102). Furthermore, we uncovered *NRXN1*, a gene encoding a membrane protein that belongs to the neurexin family, as the MAG of pathways in top communities associated with synapse organization, stimulus responses, and neurotransmitter secretion (Figure 4E). This finding aligns with its known function in the formation of synaptic contacts and efficient neurotransmission (102-104). Furthermore, the cell type enrichment and the active pathways identified in ExN showed a high overlap across multiple PFC datasets (Figure 4F-G and Supplementary Table S11). Specifically, over 93% of the active pathways identified in dataset 69 were also present in the other three datasets, indicating the robustness of our model.

In the case of PD (11-13), we focused on the substantia nigra, the brain region that is most affected by PD (105,106). We found that dopaminergic neurons (Da) showed the highest activation by PD risk genes (Figure 4H), consistent with prior knowledge of PD pathology (105,106). The top communities of Da were linked to topics related to the generation of energy, cell respiration, cell death, cell activation, chemokine-mediated signaling pathways, and vesicle and membrane organization (Figure 4I). These topics correspond to impaired mitochondrial homeostasis and defective cellular respiration reported as the factors that cause Da loss in PD (107,108). Moreover, *SNCA*, the gene encoding alpha-synuclein, was identified as the MAG of pathways in multiple top communities (Figure 4I), aligning with the known role of alpha-synuclein in PD, where its accumulation in Da leads to mitochondrial dysfunction and cell death (107,109).

Taken together, our results demonstrated CellGO’s ability to unveil cell type-specific pathogenesis of neurological disorders, highlighting its potential in studying other complex polygenic diseases.

### Web interface of CellGO

We built a website to provide an interactive interface for using CellGO without registration (Figure 5). This web interface currently provides access to pre-trained models from 71 human and mouse brain single-cell datasets and offers four analysis modes. The basic gene-set analysis mode reports cell type-specific *P*-values for pathways and cell type-specific GPPASs for genes, along with a bar plot indicating the number of active pathways per cell type. The complete gene-set analysis mode enhances the basic mode by performing pathway network analysis, providing visualizations for the network of cell type-specific pathway communities (Supplementary Figure S6) and detailed information on each community. The cell type enrichment mode calculates *P*-values for cell types based on a set of gene names, allowing for a quick assessment of cell type enrichment. The single-gene annotation mode reports cell type-specific GTSs and *P*-values for pathways directly connected to the queried gene.

**Figure 5.**
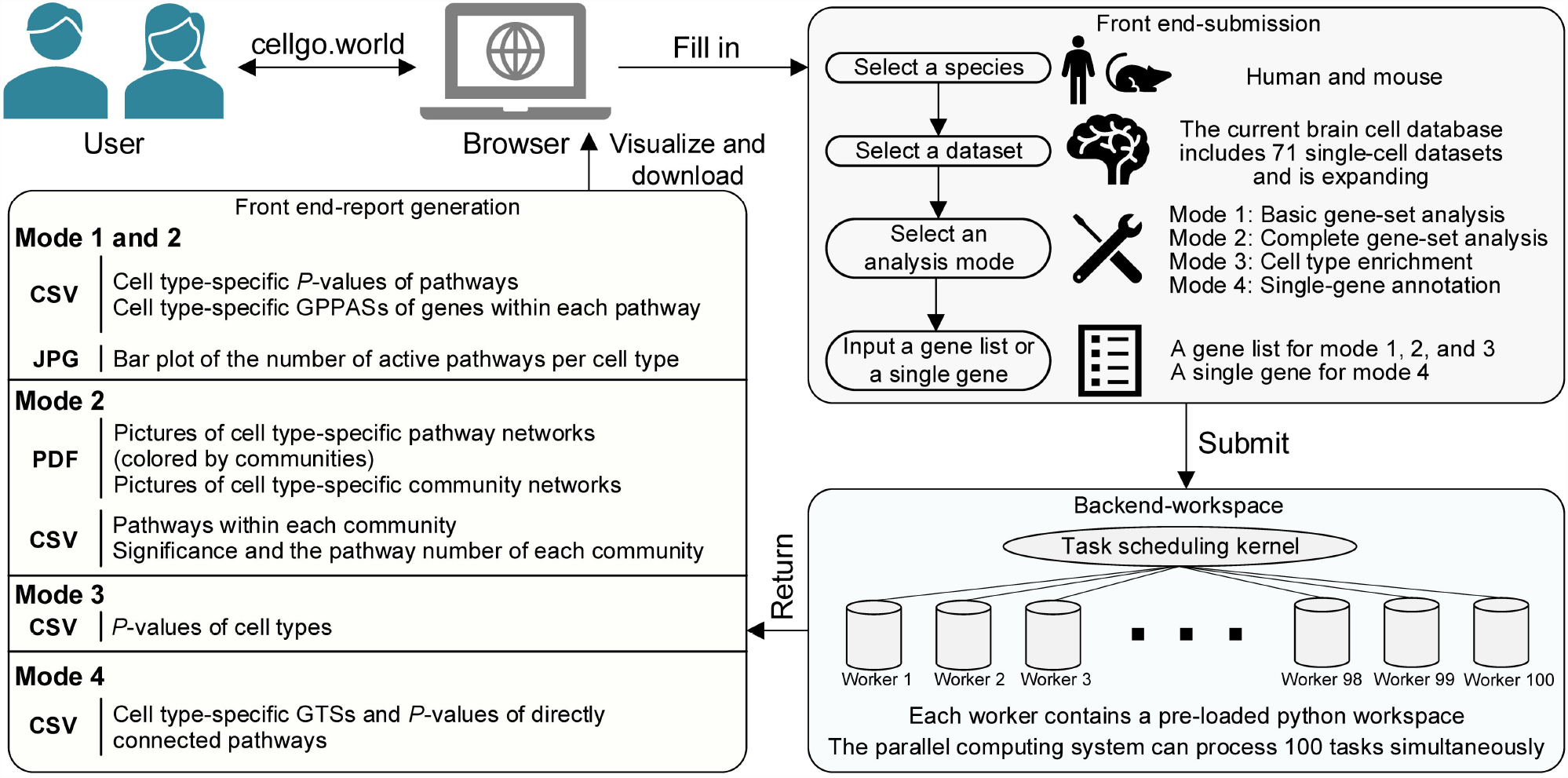
CellGO web interface. Pipeline illustration of the CellGO web interface. Users start by inputting the necessary parameters to submit an analysis request (upper right). The submission is then processed by the parallel computing system (lower right), and the results are subsequently delivered to the front end for visualization and download (left).

The web interface utilizes a task scheduling kernel to manage 100 processing-ready workers, enabling computation of up to 100 simultaneous submissions. Additionally, we encourage users to share and submit CSV files derived from the modeling phase to expand the CellGO website collaboratively with other single-cell datasets.

## 4 Discussion

Pathway analysis is a crucial step in the multi-omics workflow, offering valuable insights for mechanistic exploration (21,22). CellGO, with its unique features, represents a functionally novel tool for cell type-specific pathway analysis that takes into account the biological context of pathway activation. By simulating the cell type-specific signal propagation of genes through the hierarchical structure of the Gene Ontology tree, CellGO can predict cell type-specific pathways activated by a single gene and provide a comprehensive hierarchy of biological insights for a gene set. This innovative design significantly expands the capabilities of CellGO to provide more detailed and meaningful interpretations of complex molecular mechanisms.

The foundation of CellGO lies in its ability to consider complex interactions across genes and pathways under specific tissues and cell types during the modeling phase, which is made possible by the integration of the single-cell RNA expression data and the VNN that mirrors the functional organization of a cell. This design enables a holistic scoring system that measures the pathway activity influenced by single genes in specific cell types, and thus empowers GO annotations (40,41) with cell type-specific information, serving as the basis for the development of cell type-specific pathway analysis tools.

In addition to pathway-level significance, CellGO provides community-level significance that considers the cooperative contribution of topologically adjacent pathways in the GO graph to capture biological relevance by reducing the redundancy of synonymous pathways. Moreover, CellGO supports the inference of highly active genes within key pathways, providing additional insights into molecular origins. These advantages together broaden the ability of CellGO for mechanistic interpretation.

Our results demonstrated the effectiveness of CellGO in deciphering cell type-specific pathways influenced by gene knockout or chemical treatment. Compared to the ranked ORA-based scMappR, CellGO exhibits higher cell type specificity in pathway analysis, likely due to the adoption of a classification algorithm that tends to select cell type-specific genes and their associated pathway lineages in the VNN.

To circumvent batch effects, our website provides models pre-trained on each single-cell dataset individually. Advancements in batch correction methods may overcome this limitation in the future, allowing CellGO to be trained using combined data from multiple resources and expanding its applicability to cross-organ analysis. Another limitation of CellGO is its current inability to incorporate gene-level statistics, such as fold changes from differential expression analysis. Future versions of CellGO can address this limitation by considering fold changes during the gene expression perturbation in the modeling phase, rather than simply setting gene expression to zero.

In summary, we presented CellGO as an innovative and reliable deep learning-based tool for cell type-specific pathway analysis. We believe that it broadens the scope of pathway analysis, providing researchers with biological context-specific and multidimensional perspectives that enable them to glean valuable insights into underlying mechanisms and design further experiments.

## 5 Data availability

URLs for downloading 71 processed scRNA-seq datasets originally published by 29 studies are reported in the Supplementary Table S1. 71 re-processed scRNA-seq datasets with cell labels that organize our currently proposed brain cell database are available at http://www.cellgo.world. The bulk expression matrix for the Smith et al. *Atp1a2* KO dataset was downloaded from https://www.ncbi.nlm.nih.gov/geo/query/acc.cgi?acc=GSE145102. The bulk expression matrix for the Runge et al. *Neurod2* KO dataset was downloaded from https://www.ncbi.nlm.nih.gov/geo/query/acc.cgi?acc=GSE110491. The bulk expression matrix for the Wang et al. tau fibril dataset was downloaded from https://www.ncbi.nlm.nih.gov/geo/query/acc.cgi?acc=GSE198013.

## 6 Code availability

All the functions of CellGO have been integrated into a python package cellgopy with detailed built-in help documentation that is available at https://github.com/FduZhuLab/cellgopy. A Zenodo repository of the software is available here: https://doi.org/10.5281/zenodo.8078074. We also provide a tutorial for installation and operation of this python package at http://www.cellgo.world. The web interface for interactive execution of CellGO analysis is available at http://www.cellgo.world.

## 7 Author contributions

Y.Z. and P.L. designed the study and developed the computational methodologies. P.L. developed the cellgopy python package, and conducted computational analyses and biological interpretation with Y.Z.’s supervision. J.W. and P.L. developed the web interface. P.L. and Y.Z. wrote the manuscript with approval of all authors.

