## Supplemental Information for "CellGO: A novel deep learning-based framework and webserver for cell type-specific gene function interpretation"

### Supplementary Text

##### Application of CellGO on more gene knockout datasets

We used CellGO to analyze the differentially expressed genes generated by the other two experimental datasets (1,2). For the top 100 differentially expressed genes between *Adnp* KO and WT mice (1), CellGO found that microglia had the highest activation according to the number of active pathways (Supplementary Figure S4B). Noteworthily, top communities of microglia represented topics related to metabolic and immune system processes, stimulus responses, signal transduction, and reproductive processes (Supplementary Table S8). More specifically, organic substance metabolic process, immune response-inhibiting signal transduction, positive regulation of leukocyte differentiation, and reproductive process were found in these communities (Supplementary Figure S4B), with *Hexb* (a gene encoding the beta subunit of the lysosomal enzyme beta-hexosaminidase that breaks down fatty compounds and complex sugars), *Lyn* (a gene encoding the tyrosine protein kinase that involves in immune response-regulating signaling pathways), and *Inpp5d* (a gene encoding the protein that functions as a negative regulator of myeloid cell proliferation and survival) identified as the MAGs in these pathways. These findings are consistent with previous studies (1) showing that *Adnp* KO leads to pro-inflammatory and pro-phagocytic phenotype and a prominent increased number of microglia.

For the top 100 differentially expressed genes between neuron-specific *Wwox* knockdown and WT mice (2), CellGO discovered that oligodendrocytes showed the highest activation (Supplementary Figure S4C). Notably, top communities of oligodendrocytes represented topics related to cell development, metabolic processes, homeostatic processes, signaling, and ion transport (Supplementary Table S8). More concretely, glial cell development, cell differentiation, transmission of nerve impulse, and positive regulation of calcium ion transmembrane transport were found in these communities (Supplementary Figure S4C), with *Plp1* (a gene encoding a transmembrane proteolipid protein that is the predominant component of myelin and plays a role in oligodendrocyte development and axonal survival) and *Nfasc* (a gene encoding the protein functions in neurite outgrowth, neurite fasciculation, and organization of the axon initial segment) predicted as the MAGs in these pathways. These findings align with a previous study (2) indicating that *Adnp* knockdown in neurons reduced oligodendrocyte maturation and induced hypomyelination and impaired axonal conductivity. These results again demonstrated that CellGO could be used effectively to infer functionally relevant cell types and cell type-specific pathways.

### Supplementary Figure

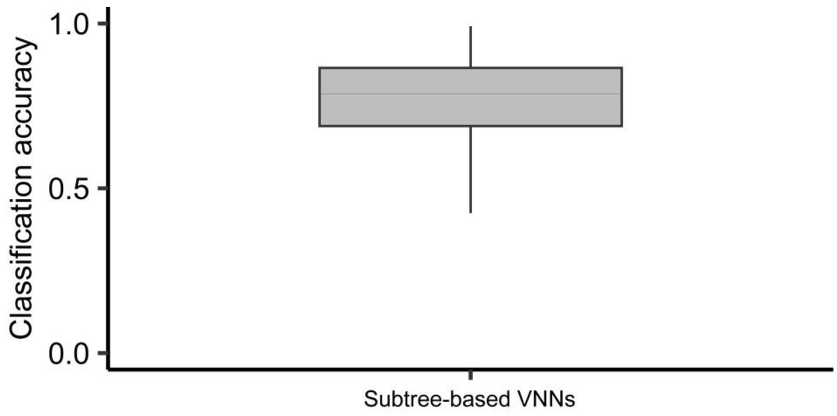


**Supplementary Figure 1.** The accuracy of CellGO's performance in cell type classification.
Boxplot indicating the classification accuracy of 4,168 subtree-based VNNs using dataset 1. The boxplot displays the median as the horizontal middle line, the 0.25 quantile ($Q_1$) and 0.75 quantile ($Q_3$) as the bounds of the box, and the upper and lower whiskers based on the minimum of the maximum value and $Q_3 + 1.5*IQR$, and the maximum of the minimum value and $Q_1 - 1.5*IQR$, respectively.

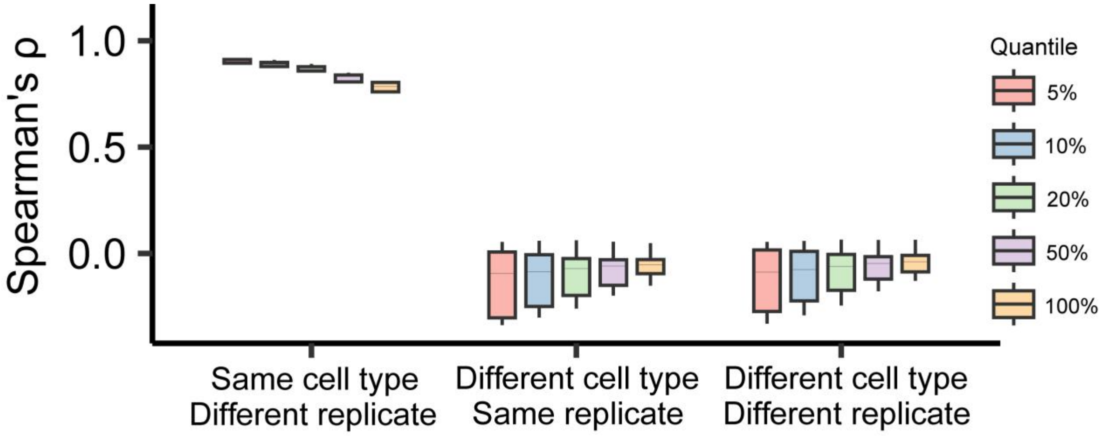


**Supplementary Figure 2.** Assessment of model reproducibility.
Boxplot showing pairwise Spearman's rank correlation coefficients across GTSs of the same cell type from different replicates (left), GTSs of different cell types from the same replicate (middle), and GTSs of different cell types from different replicates (right), calculated using four independent replicates of dataset 1. Spearman's correlations calculated using different quantiles of gene-term pairs were displayed. For example, 5% indicates that the correlations were calculated on gene-term pairs with GTSs ranked in the top 5% in either group. The boxplot displays the median as the horizontal middle line, the 0.25 quantile ($Q_1$) and 0.75 quantile ($Q_3$) as the bounds of the box, and the upper and lower whiskers based on the minimum of the maximum value and $Q_3 + 1.5*IQR$, and the maximum of the minimum value and $Q_1 - 1.5*IQR$, respectively.

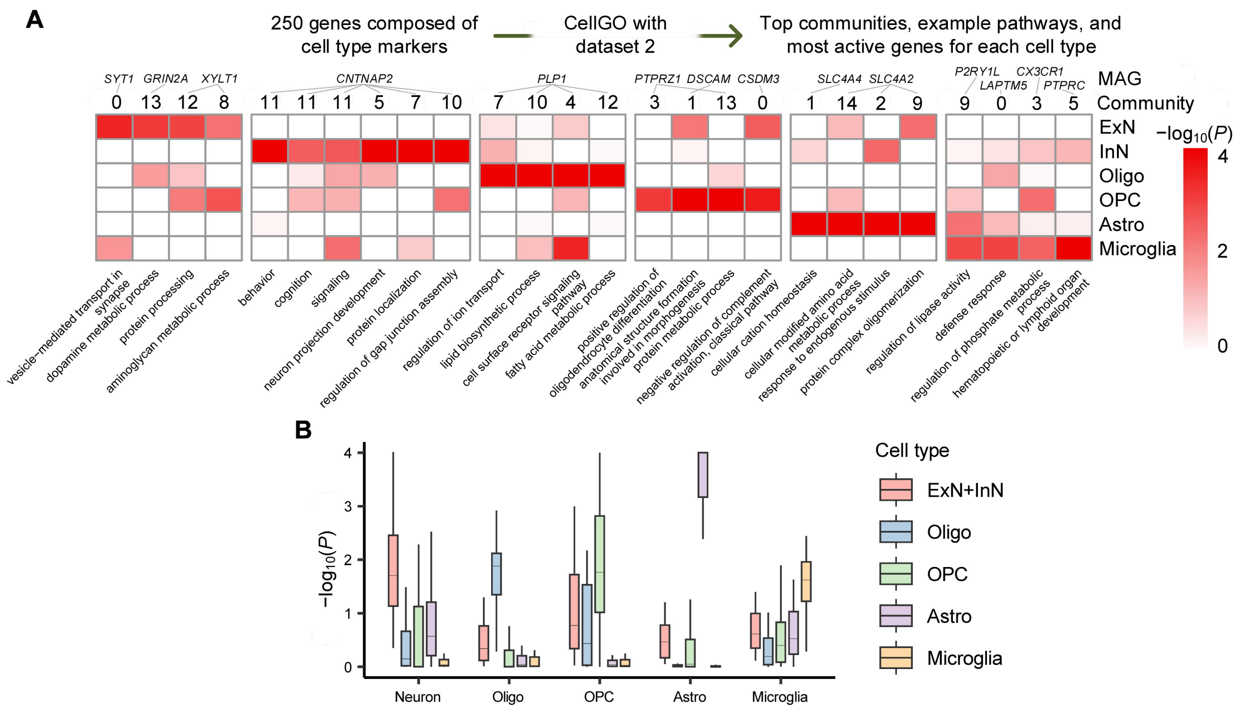


**Supplementary Figure 3.** Application of CellGO to capture cell type-specific signatures using dataset 2. (**A**) Top four communities, example pathways in these communities, and MAGs of these pathways for each cell type identified by CellGO testing 250 cell type markers using dataset 2. Heatmap colors denote the cell type-specific log-transformed *P*-values of example pathways in the top four communities per cell type. The community index and the MAG of each pathway are shown above the heatmap.
(**B**) Box plot showing cell type-specific log-transformed *P*-values from CellGO for predefined cell type-specific pathways. The boxplot displays the median as the horizontal middle line, the 0.25 quantile ($Q_1$) and 0.75 quantile ($Q_3$) as the bounds of the box, and the upper and lower whiskers based on the minimum of the maximum value and $Q_3 + 1.5*IQR$, and the maximum of the minimum value and $Q_1 - 1.5*IQR$, respectively.

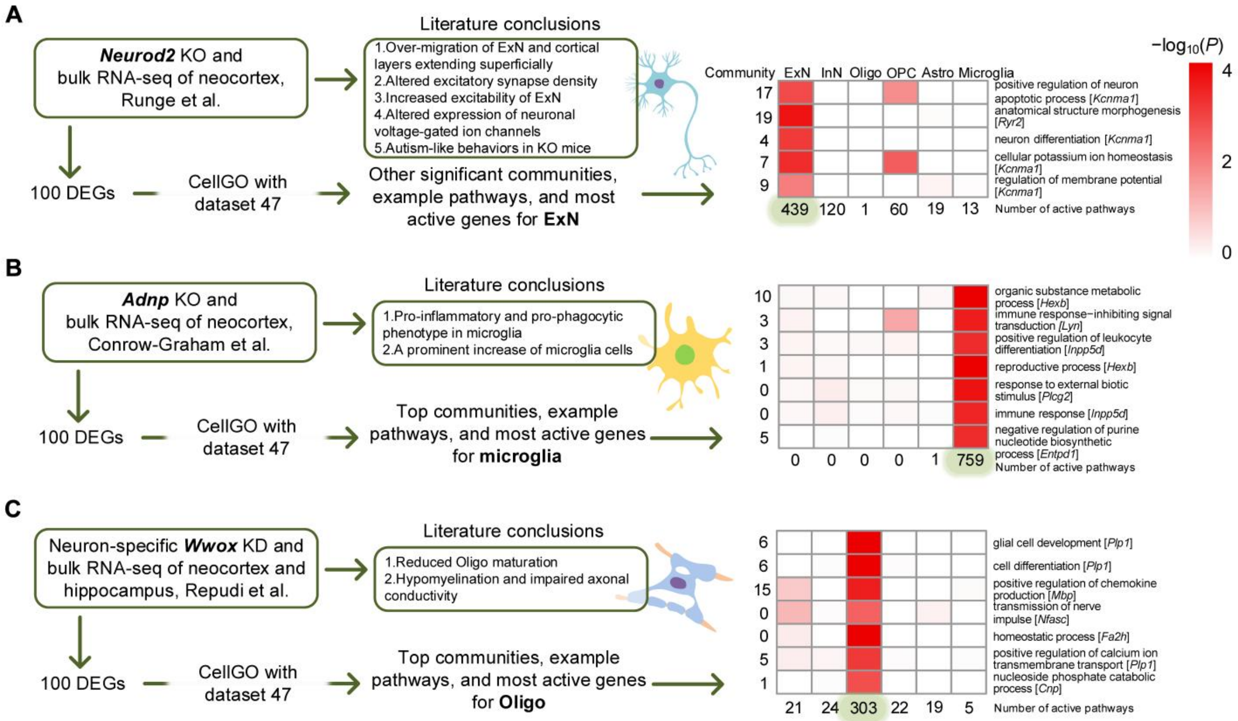


**Supplementary Figure 4.** Benchmarking of CellGO's performance on more experimental datasets. (**A,B,C**), Results obtained from CellGO when analyzing differentially expressed gene lists from (**A**) Runge et al., (**B**) Conrow-Graham et al., and (**C**) Repudi et al.. The main conclusions reported in the original studies are listed as ground truth. Heatmap colors represent log-transformed *P*-values. (**A**) More significantly active pathways not in the top five communities associated with *Neurod2* are presented. (**B,C**) Example pathways in the top five communities of the most active cell type are plotted. The number of active pathways (*P*-value < 0.01) per cell type is displayed underneath the heatmap, and the MAG of each pathway is provided next to the pathway name.

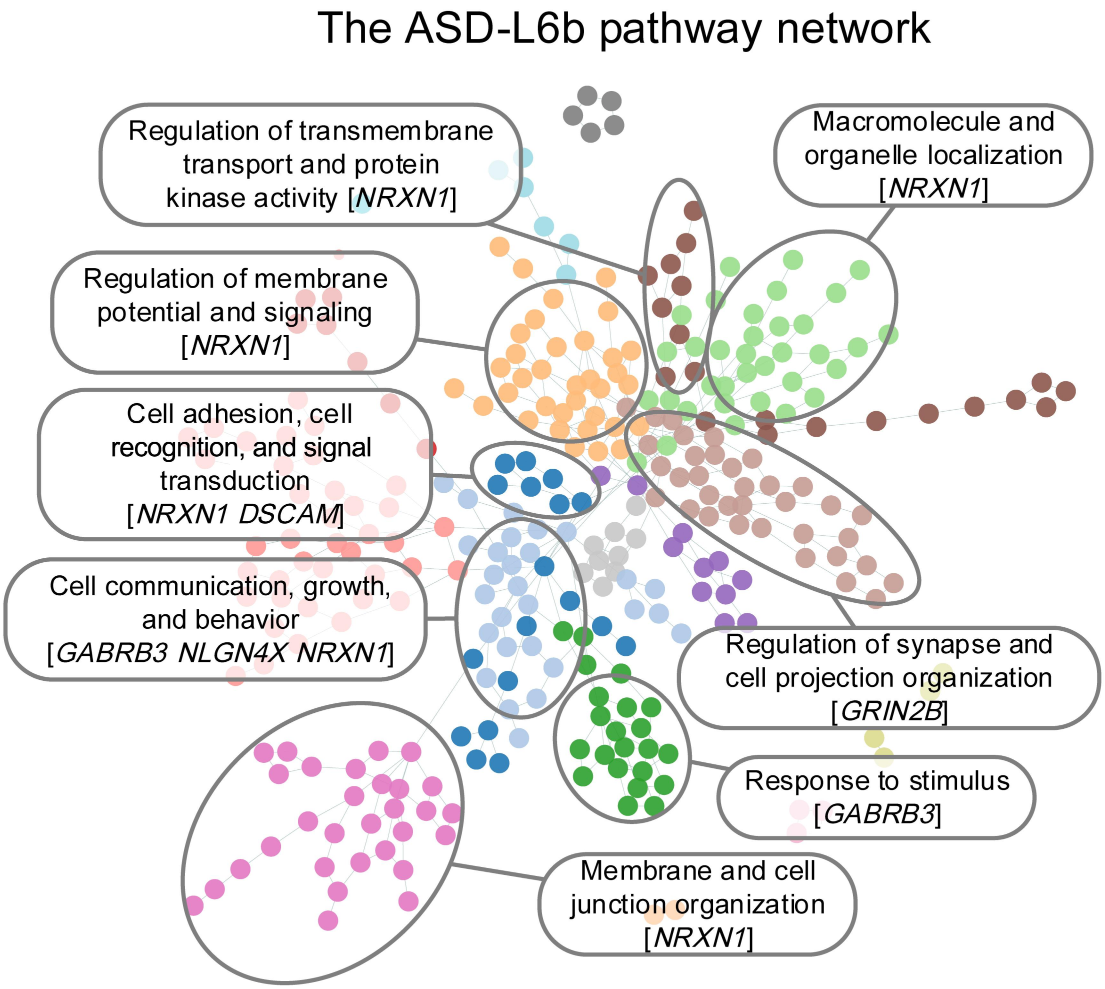


**Supplementary Figure 5.** The ASD-associated L6b-specific pathway network.
The pathways are colored by communities, with the top eight communities elliptically labeled and annotated with topics. MAGs of pathways with close semantics to the topic of communities are labeled.

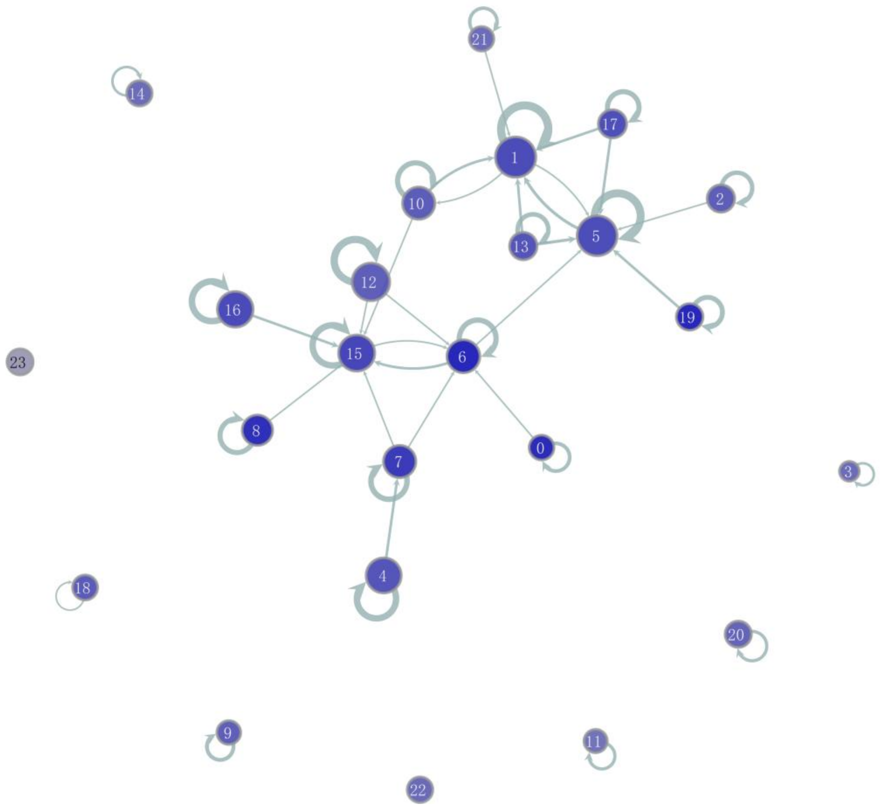


**Supplementary Figure 6.** The ASD-associated ExN-specific community network.
This community network is matched with the Fig. 4e. Each node represents a pathway community, and the community number is shown in the node; the color of each node denotes the significance of the community; the size of each node denotes the number of pathways in the community; the thickness of each edge between nodes denotes the number of connections between communities.

### Description of Supplementary Tables

**Supplementary Table 1**
Detailed information of the curated brain cell database
**Supplementary Table 2**
Cell type markers
**Supplementary Table 3**
Predefined cell type-specific pathways
**Supplementary Table 4**
Five experimental datasets and corresponding DEG lists for testing
**Supplementary Table 5**
Outputs from CellGO analyzing cell-type marker genes based on dataset 1
**Supplementary Table 6**
Outputs from CellGO analyzing cell-type marker genes based on dataset 2
**Supplementary Table 7**
CellGO outputs form single-gene-mode analysis of *Atp1a2* and *Neurod2*
**Supplementary Table 8**
CellGO, gProfileR, and scMappR outputs of the five DEG lists
**Supplementary Table 9**
Disease risk genes
**Supplementary Table 10**
CellGO outputs of disease risk genes
**Supplementary Table 11**
CellGO outputs of ASD risk genes based on four PFC datasets
